# Biomechanical Stimulation of Muscles Influences Bone Phenotype by Modulating Myokine Secretion

**DOI:** 10.1101/2022.10.05.510953

**Authors:** Harshini Suresh Kumar, Edwina N. Barnett, Evangelia Kalaitzoglou, John L. Fowlkes, Ramkumar T. Annamalai

**Author notes:** **Author Correspondence:** Ramkumar T. Annamalai, 760 Press Avenue, 138 Healthy Kentucky Research Building, University of Kentucky, Lexington, KY 40536, **Email:**.

## Abstract

Diabetes is a chronic metabolic disorder that affects 422 million people worldwide and can lead to diabetic myopathy and bone diseases. The etiology of musculoskeletal complications in diabetes and the interplay between the muscular and osseous systems are poorly understood. Exercise training promises to prevent diabetic myopathy and diabetic bone disease and offer protective effects on muscle and bone. Although the muscle-bone interaction is largely biomechanical, the muscle secretome, specifically the myokines, has significant implications for bone biology. Here, we have developed an *in vitro* model to elucidate the effects of mechanical strain on myokine secretion and its impact on bone metabolism decoupled from physical stimuli. We developed modular bone constructs using crosslinked gelatin hydrogels which facilitated osteogenic differentiation of osteoprogenitor cells. Then muscle constructs were made from fibrin hydrogel, which enabled myoblast differentiation and formed mature myotubes. We investigated the myokine expression by the muscle constructs under strain regimens replicating endurance (END) and high-intensity interval training (HIIT) in hyperglycemic conditions. In monocultures, both regimens induced higher expression of *Il15* and *Igf1*, while END supported more myoblasts differentiation and myotube maturation than HIIT. When cocultured with bone constructs, the HIIT regimen increased *Glut4* expression in muscle contructs that END supporting higher glucose uptake. Likewise, the muscle constructs under the HIIT regimen promoted a healthier and matured bone phenotype than END. Interestingly, under static conditions, myostatin (*Mstn)* expression was significantly downregulated in muscle constructs cocultured with bone constructs compared to monocultures. Our *in vivo* analysis of the role of myostatin on bone structure and function also showed that myostatin knockout (GDF8^-/-^) enhanced muscle mass and moderately influenced bone phenotype in adult mice. Together, our *in vitro* coculture system allowed orthogonal manipulation of mechanical strain on muscle constructs while facilitating biochemical crosstalk between bone and muscle constructs. Such systems can provide an individualized microenvironment and allow decoupled biomechanical manipulation, which is unachievable using traditional models. In the long-term, these in-vitro systems will help identify molecular targets and develop engineered therapies for diabetic bone disease.

## 1. Introduction

Diabetes mellitus is a collection of metabolic illnesses defined by long-term hyperglycemia caused by defects in insulin secretion and action^1-3^. According to the World Health Organization, 422 million people worldwide suffer from diabetes mellitus, and over 11.3% of the US population has been diagnosed with diabetes (Centre for Disease Control and Prevention, US, 2021). Diabetes-related complications negatively impact the quality of life and can lead to financial hardships. Diabetes can affect every organ system in the human body, and the extent of pathophysiology depends on the severity and duration of the disease. Microvascular complications such as retinopathy, nephropathy, and neuropathy, as well as macrovascular complications such as cardiovascular disease, are the most commonly recognized diabetic-associated complications^4^. Diabetes also leads to major musculoskeletal complications, including diabetic myopathy and diabetic bone disease. Diabetic myopathy is characterized by skeletal muscle loss and impaired function secondary to muscular inflammation, ischemia, hemorrhage, infarction, necrosis, fibrosis, and fatty atrophy^5^. Diabetic bone disease is characterized by poor bone quality, leading to increased fracture risk, osteoporosis, and delayed bone healing^6^. Several factors such as severity and duration of diabetes, age at diagnosis, lack of resistance to endogenous insulin and micro-and macrovascular diabetes complications have been implicated in the development of diabetic myopathy^7, 8^ and diabetic bone disease^9, 10^. Additionally, treatment of diabetes with oral or injectable hypoglycemic agents can have negative effects on the musculoskeletal system^10, 11^; for example thiazolidinediones have been associated with bone loss and increased risk of fracture^12^ and DPP4 inhibitors have been shown to cause myalgia, muscle weakness and other side effects in skeletal muscle^13^. Several prevalent musculoskeletal disorders, including muscle cramps and carpal tunnel syndrome, are now recognized as a natural result of elevated glucose levels^14^. However, the underlying mechanisms of these musculoskeletal complications in diabetes and the interplay between the muscular and osseous systems are poorly understood.

For diabetes-mediated musculoskeletal pathologies such as myopathy, symptomatic alleviation is the primary treatment that involves pain management, rest, and anti-inflammatory medicines^15^. Pharmacotherapies using disease-modifying agents such as anti-myostatin peptides, *STAT3* inhibitors, anabolic androgenic steroids, *IGF-1*, and antisense oligonucleotides have been studied to treat diabetic myopathy^16^. Likewise, antiresorptive or anabolic medications containing bisphosphonates could be useful in preventing diabetes-related fractures. Though these pharmaceutical agents effectively ameliorate specific symptoms, they often cause off-target effects. Physical exercise has shown beneficial effects, including improved glucose metabolism and insulin sensitivity, and is often recommended along with pharmacotherapies for the clinical management of diabetes^17^. An acute bout of exercise increases muscle glucose uptake efficiency, and chronic exercise training enhances mitochondrial biogenesis, increasing the expression levels of glucose transporter proteins and various metabolic genes^18^. In addition, these muscular contractions can promote healthy metabolism in other tissues and organs, including bone. For decades, researchers have investigated muscle contraction-induced ‘exercise factors’ that mediate muscle-bone synergism.

The muscle-bone interaction is primarily mechanical, with muscle imposing contractile forces on bone tissue^19^. But skeletal muscle also acts as an endocrine organ by secreting factors called myokines that are implicated in various physiological and pathological mechanisms in other organs, including bone^20-22^. Factors related to exercise training such as duration, intensity, muscle mass, and endurance capacity determine the type and magnitude of myokine secretion from skeletal muscle^23, 24^. A subset of myokines produced by skeletal muscle contraction can positively influence bone formation and regulate glucose uptake rate. Under hyperglycemic conditions, the synthesis and secretion of several myokines are altered^25^. It has been hypothesized that this diabetic-mediated imbalance in myokine secretion plays a crucial role in developing diabetic myopathy and bone diseases^26, 27^. Myostatin and irisin are key myokines known to promote osteoclastogenesis and osteoblast differentiation, respectively^28, 29^. Although myostatin is critical for osteoclast development, it can have detrimental effects on bone mass by reducing bone formation and increasing bone resorption^30^. Remarkably, weight-bearing exercises are shown to inhibit myostatin expression and reduce the blood levels of alkaline phosphatase (ALP) and tartrate-resistant acid phosphatase (TRAP) in a rat T1D model^31^. On the other hand, irisin can upregulate osteogenic genes such as *RUNX2*, Osterix (*OSX*), *ATF4*, β-catenin (*CTNNB1*), alkaline phosphatase (*ALP*), and type I collagen (*COL1A1*) in osteopotent cells. Follistatin, an endogenous inhibitor of myostatin, is shown to increase after resistance exercise. Resistance exercise activates mTORC1 mediated increase in miR-1, which decreases myostatin activity by secreting follistatin^32^. Some anti-inflammatory cytokines, including IL-10, IL-1Ra, and soluble receptors of the tumor necrosis factor (TNF) I and II, were also shown to increase following acute aerobic exercise^33^. However, the underlying cellular pathways regulating the muscle secretome during exercise training and its protective effects on bone physiology and function are poorly understood. Hence it is crucial to understand the effect of exercise on myokine secretion pattern and its effect on the bone to determine molecular targets for developing therapies for diabetes-mediated myopathy and bone disease.

Uncoupling the effects of biophysical and biochemical stimuli on the adaptive response of bone during exercise training is challenging using animal models. Here, we have developed an *in vitro* culture system to elucidate the effects of mechanical strain on myokine secretion and its impact on bone metabolism decoupled from physical stimuli. We fabricated bone and muscle constructs using natural polymers and configurations that allow the cells to exhibit their innate tissue-specific phenotype. We used crosslinked gelatin microspheres and seeded them with preosteoblasts to fabricate bone constructs, as shown previously^34, 35^. Gelatin is obtained from partially hydrolyzed collagen retaining RGD motifs for cell attachment, biocompatibility, and osteoconductivity^36^. In addition, the microsphere format provides a 3D microenvironment with physical cues that can promote osteogenic differentiation of progenitor cells^37^. Likewise, the muscle constructs were fabricated by seeding myoblasts in 3D fibrin hydrogels. Fibrin facilitates muscle cell migration and proliferation without spatial inhibition enabling them to self-organize into myotubes^38^ and has been used to regenerate skeletal muscles and treat muscular dystrophies. Further, fibrin is known to bind to angiogenic factors, including vascular endothelial growth factor (VEGF), basic fibroblast growth factor -2 (bFGF-2), and insulin-like growth factor -1 (IGF) that participate in myogenesis^39^. Together, our system provides an individualized microenvironment and allows decoupled biomechanical manipulation, which is not achievable in animal models. Then we developed an *in vitro* culture apparatus using these engineered constructs that allowed orthogonal manipulation of mechanical strain on muscle constructs while facilitating biochemical crosstalk between bone and muscle constructs, as shown in **Fig.1**. We hypothesize that the type and intensity of mechanical strain regulate the myokine secretion, which influences myotube and osteoblast phenotype and function in a hyperglycemic microenvironment. We investigated the secretion of myotubes under endurance (END) and high-intensity interval (HIIT) strain regimens under hyperglycemic conditions and their effects on bone metabolism and vice versa. The effect of myostatin, a key myokine implicated in several diabetes-driven pathologies, on bone constructs was investigated. The function of follistatin to revert the adverse effect of myostatin on bone cells was also explored. We also investigated the effect of myostatin inhibition on bone anatomy and physiology in mice by characterizing their mechanical and material properties. Overall, our studies elucidate the effects of biophysical and biochemical stimuli on muscle secretome and its corresponding impact on bone physiology and vice versa in a hyperglycemic environment. Our work is crucial and can help identify therapeutic targets and develop engineered therapies for various diabetic-related complications.

**Fig 1.**
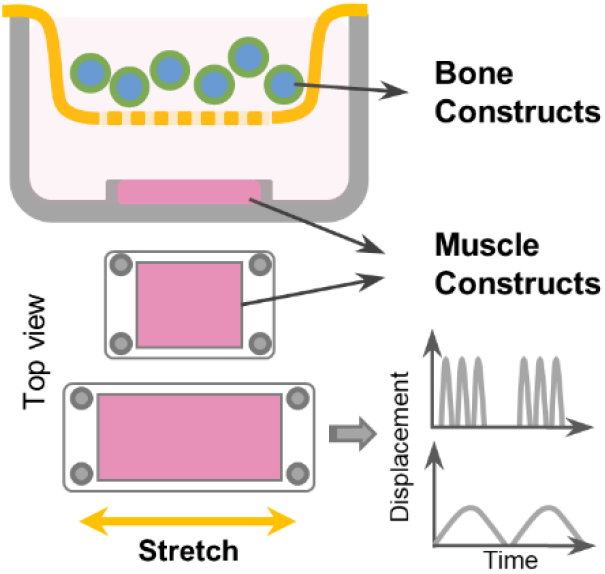
Muscle-bone crosstalk in diabetes. Coculture system to elucidate the impact of muscle exercise on bone metabolism.

## 2. Materials and Methods

### 2.1. Fabrication of Bone constructs

To fabricate bone constructs, we made 3D microgels (∼150 µm dia.) from solubilized gelatin through a simple water-in-oil emulsification process and crosslinked them using genipin (Wako), as described^34, 35^. Briefly, the 6 wt.% gelatin solution (Porcine skin, Type A, Sigma) in deionized (DI) water was added to a beaker containing stirred polydimethylsiloxane (PDMS) at 37°C and emulsified for 5 minutes. The mixture was stirred at 450 rpm using a spiral propeller blade, and the emulsion was then cooled to 4°C for 30 min using an ice bath. The emulsion was then centrifuged at 175 g for 5 min to separate the gelatin microgels from PDMS. The pellet was washed thrice with 10 mM PBS (Gibco) containing 1% TWEEN 20 (Sigma, PBS-T20) and suspended in genipin (1 wt.% in 1X PBS) for 48 hours. The crosslinked microgels were then washed in 100% ethanol to remove excess genipin. The microgels were filtered using nylon sieves to obtain the desired size range of 100 – 150 µm in diameter. To fabricate bone constructs, the microgels were sterilized with ethanol overnight and seeded with murine preosteoblast cell line (MC3T3, Riken-BRC) at a density of 4×10^6^ cells/mg dry mass of microgels. The cells were cultured in α Modified Eagle’s Medium (α-MEM, Gibco) supplemented with 10% v/v FBS, 1X antibiotic-antimycotic solution (AA, Gibco), and 0.2% normocin (Invivogen) and maintained in a humidified CO_2_ incubator. To induce osteogenic differentiation, α-MEM supplemented with 10% FBS, 1% 20mM Ascorbic Acid 2 Phosphate, 1% 1M β glycerol phosphate was added to the microgel suspension and cultured for 5 days.

### 2.2. Fabrication of Muscle Constructs

To fabricate muscle constructs, we used murine myoblasts (C2C12, ATCC) encapsulated in fibrin hydrogel. To make the fibrin hydrogel, fibrinogen stock (4 mg of clottable protein/mL of DMEM, bovine plasma, Type I-S, Sigma) was filter-sterilized and mixed with myoblasts suspension. Thrombin (Sigma) was added to cleave the fibrinogen molecule and form 3D fibrin gel. The final concentration of the mixture was 2.5 mg/mL of fibrinogen/fibrin, 1 U/mL of thrombin, and a cell density of 2×10^6^ cells/mL. The mixture was then quickly cast into specially molded 12×12 mm PDMS wells and placed in a humidified incubator for 45 min to allow complete gelation. The cells were supplied with growth media (DMEM-GlutaMax (4.5 g/L of glucose, 10% FBS, 1X AA, and 0.2% normocin) for 48 hours and switched to differentiation media (DMEM-GlutaMax) supplemented with 2% horse serum (Sigma), 1% AA, and 0.2% normocin) and differentiated for 5 days. Media was replenished daily till the mature myotubes formed were verified. Fully matured muscle constructs were used for characterization and coculture experiments.

### 2.3. Exercise Training of Muscle Constructs

The muscle constructs were subjected to linear strain using Cytostretcher™ (CuriBio, Seattle, WA). Muscle constructs with mature myotubes were subjected to acute high-intensity strain (10% strain, 1 Hz, 4 hours/day in bouts of 1 hour) or chronic moderate strain (1%, 0.1 Hz, 12 hours/day) mimicking high-intensity interval training (HIIT) and endurance (END) exercise, respectively. Static cultures served as a control. Media samples were collected at specific time points and analyzed for various myokine secretions. For the coculture experiments, the bone constructs were kept above the muscle constructs either using membrane-bottom rectangular trans wells (5 × 5 mm) or directly on top of the muscle constructs to allow biochemical communications between the constructs without any strain transfer.

### 2.4. Immunofluorescence and Confocal microscopy

Bone and muscle samples collected at different time points were fixed in 10% buffered formalin (Thermo Scientific), permeabilized with 0.1% Triton X-100 (Sigma) in 1X PBS, and blocked with 1% bovine serum albumin (BSA) in 10 mM PBS. The cells were then stained for cytoskeletal F-actin using phalloidin conjugated with Alexa fluor 488 or Alexa fluor 594 (Invitrogen) and DAPI (Life Technologies) as nuclear counterstaining. Additionally, the muscle construct was stained for myosin heavy chain to characterize myotube formation. The muscle construct was incubated with the primary antibody for myosin heavy chain (Sigma M4276, 1:400 dilution) for 2 hours at room temperature. Then the samples were washed thrice in PBS-T20, and a goat anti-mouse IgG1 secondary antibody conjugated with Alexa fluor 488 (Invitrogen A21121, 1:500) was added and incubated for 1 hour at room temperature. Then the samples were mounted in ProLong™ Gold anti-fade media (for 2D cultures only), and confocal stacks were acquired using Nikon A1R inverted confocal microscope.

### 2.5. Electron microscopy

Bone and muscle samples collected at different time points were fixed with 3% glutaraldehyde for a minimum of 1 hour. Constructs were then washed in PBS and DI water and suspended in 1% osmium tetroxide for 16 hours at 4°C. Then the samples were washed in DI water, dehydrated using an ethanol series (30%, 50%, 70%, 80%, 90%, 96%, 100%), and freeze-dried overnight. The dried samples were then mounted to stubs containing carbon adhesive, coated with 5 nm layer of platinum using a sputter coater (Leica ACE600). Then the coated samples were imaged under a scanning electron microscope (FEI Quanta 250).

### 2.6. Compression Testing

The stiffness of the genipin crosslinked gelatin hydrogel was determined using the compression testing of cylindrical blocks of the samples using a universal mechanical testing machine (Instron 6800). Briefly, the gelatin-genipin hydrogel was cast between two parallel glass plates separated by a 5 mm spacer. Using an 8 mm biopsy punch, samples were punched out and compressed at a rate of 2 mm/min to obtain the stress-strain curve. The young’s modulus was then calculated from the slope of the linear portion of the curve between 5-15% strain.

### 2.7. Myostatin and Follistatin treatment

The preosteoblast cells cultured in tissue culture plates were differentiated for 5 days and treated with Myostatin of 100 ng/mL (788-G8-010/CF R&D system), follistatin of 100 ng/mL (769-FS-025/CF, R&D system) and a cocktail of myostatin and follistatin (overall 50 ng/mL each) for two or 7 days. The cells were detached and analyzed for the expression of prominent osteogenic genes.

### 2.8. Quantitative Gene Expression

The samples were collected in TRIzol reagent (Invitrogen), and the total RNA was isolated following the manufacturer’s protocol. 100 ng of RNA was used for gene expression assay. Real-time quantitative polymerase chain reaction was carried out in duplicates with a reaction volumes of 10 µL using TaqMan probes (Advanced Biosciences) for myogenic genes *Igf-1 (Insulin growth factor 1* Mm00439560_m1*), Fgf-2 (*Fibroblast growth factor Mm00433287_m1*), Mstn (*Myostatin, Mm01254559_m1*), Fndc-5 (*Irisin, Mm01181543_m1*), Il-15 (*Interleukin-15, Mm00434210_m1*), Myog (*Myogenin, Mm00446194_m1), *Myod1(*Myoblast determination protein-1 Mm00440387_m1*), Fst (*Follistatin, Mm00514982_m1*), Glut4 (*Glucose transporter type 4, Mm00436615_m1), *Ppia (*Peptidylprolyl isomerase, Mm02342430_g1) and osteogenic – *Alp1 (*Alkaline phosphatase 1, Mm00475834_m1*), Col1a1(*Collagen type I alpha 1 Mm00801666_g1*), Runx-2 (*Runt-related transcription factor 2, Mm00501584_m1*), Bglap (*Osteoclacin, Mm03413826_mH*), Spp1 (*Osteopontin, Mm00436767_m1*), Ibsp (*Integrin binding sialoprotein, Mm00492555_m1*), Sp-7, (*Osterix, Mm04209856_m1*). Gapdh (G*lyceraldehyde-3-phosphate dehydrogenase, Mm99999915_g1*), and Actb* (Actin beta, Mm02619580_g1) was used as internal controls. A SuperScript™ III Platinum™ One-Step qRT-PCR Kit (ThermoFisher) was used for reverse transcription, and the gene expression study was performed using QuantStudio 3 real-time PCR system (ThermoFisher).

### 2.9. Animal care and tissue collection

All studies involving animal cells or tissues are conducted according to approved protocols by the Institutional Biosafety Committee (IBC) protocol and the Institutional Animal Care and Use Committee (IACUC) at the University of Kentucky. Wildtype and myostatin knockout male mice of C57BL6 background and 7-10 months of age were included in this study. Myostatin knockout was induced using VelociGene™ (substituting the myostatin gene with LacZ, Regeneron)^40^. The knockout was confirmed by genotyping for 494mTD and LAcZD. All mice were housed in sterile cages and fed with a basic chow diet (#2918, Envigo). Animals were euthanized by CO_2_ asphyxiation followed by cervical dislocation. One set of femurs was fixed in 10% formalin overnight and stored in PBS for micro-computer tomography (microCT) analysis. The other set of femurs was collected in DMEM media, and a 3 point-bending test was performed within 3 hours.

### 2.10. Microcomputer tomography scanning and analysis

Femurs were imaged with a high-resolution microCT scanner (SkyScan 1276, Bruker microCT, Belgium). The regions of interest (ROIs) for the femur included the mid shaft (1.86 mm) at the point of loading. The scan parameters were: 70 kV/200 µA, 1 mm Al filter, beam hardening correction, 500 ms exposure time, and 20.38 µm image pixel size. The bone mineral density was calibrated using 0.25 and 0.75 g/cm^3^ calcium hydroxyapatite (CaHA) phantoms. The bone volume/tissue volume, bone surface/ bone volume, trabecular thickness, and total porosity was calculated using CTan software. The properties were adjusted for body mass using the linear regression method, as described^41^. CTvox software was used to construct 3D volumerendering images of the trabecular bone.

### 2.11. Biomechanical testing and analysis

Biomechanical properties were evaluated using a three-point bending test, as described previously^42^. Briefly, the width of the hydrated femurs were measured on both sides of the short axis and were loaded to failure in a three-point (3 pt) bending at a displacement rate of 3 mm/min and a support span width of 8 mm using a hydraulic material testing system (Instron 6800) fitted with a 10 N load cell. All femurs were placed on the support with femoral condyles facing upwards, and the test was performed at room temperature. The force-displacement curve was acquired and processed to determine structural properties, including stiffness or slope, peak force, yield force, post-yield displacement, and work to failure as described^42^. The stiffness was calculated as the slope of the initial linear portion of the force-displacement curve. Young’s modulus was measured as the slope of the linear portion of the stress-strain curve using the Bluehill^®^ universal software (Instron). Briefly, data on the stress-strain axis (below the yield point) between the start and end value is divided into 6 equal regions with no overlap. Then a the least square fit method was applied to all points in each region to determine the corresponding slope. The pair of consecutive regions that has the highest slope sum is determined, and the highest slope value is assigned as the modulus of the sample. For normalization, the structure and mechanical properties measured were adjusted to the animal’s body mass using the linear regression method, as described^41^. Briefly, the following equation was used to normalize unadjusted traits of individual mice with their corresponding body mass.

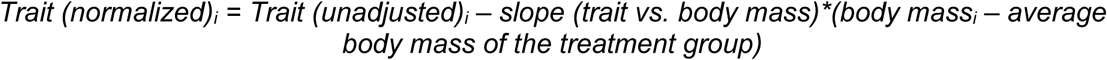

### 2.12. Statistics

All experiments were performed in at least triplicates. Statistical comparisons were made using Student’s *t*-test and one-way ANOVA with a 95% confidence limit. Dunnett test was performed to compare the treated groups with a single control group and the Tukey test to compare independent groups. Differences with p<0.05 were considered statistically significant.

## 3. Results

### 3.1. Bone constructs support osteogenic differentiation of progenitor cells

We fabricated crosslinked gelatin microgels with a size range of 150-250 µm (**Fig.2A**), sterilized them, and seeded them with preosteoblast (MC3T3 cells). The cells readily attached to the microgel surface within an hour of seeding (**Fig.2B**) and exhibited an osteoblast phenotype with rich actin cytoskeleton by 24 hours (**Fig.2C**). The progenitor cells proliferated on the surface of the microgel, and when maintained for an extended period they tended to form aggregates in suspension cultures. Using compression testing, we found that crosslinked gelatin has an elastic modulus of 111.61±12.1 kPa while the uncrosslinked hydrogels had only 24.8±3.3 kPa (**Fig.2D**). Then, the osteogenic potential of the microgels was assessed using long-term cultures. The osteoblast progenitors were seeded on microgels, maintained in growth and osteogenic conditions, collected at various time points, and analyzed for prominent osteogenic marker expression. In osteogenic culture conditions, the progenitor cells differentiated and upregulated the expression of prominent osteogenic genes within a week (**Fig.2E**). There was a significant increase in the expression of *Alp1* (Alkaline phosphatase), *Bglap* (Osteocalcin), *Ibsp* (Bone sialoprotein), and *Spp1* (Osteopontin) by day 7 (See **Table 1** for groupwise comparison). There was no significant change in osteogenic gene expression under growth conditions. These results suggest that the crosslinked gelatin constructs were conducive to osteogenesis and appropriate for fabricating bone constructs.

**Figure 2.**
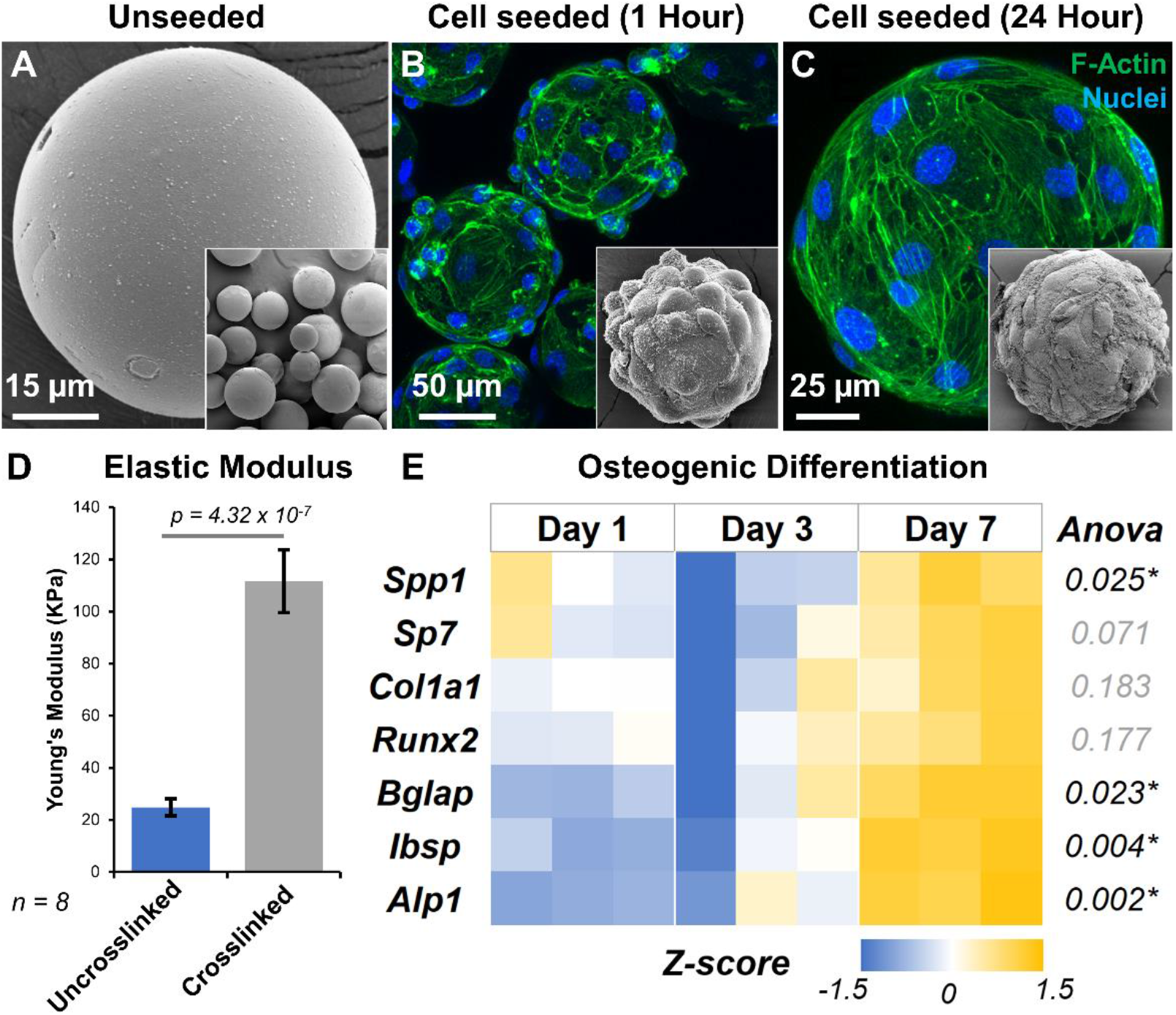
Fabrication and characterization of bone constructs. **A)** Scanning electron micrograph of lyophilized microgels showing smooth surface morphology. Confocal stacks of preosteoblasts on microgel surface **B)** 1 hour after seeding and **C)** 24 hours after seeding. The insets show electron micrographs of the corresponding samples. **D)** Elastic modulus of crosslinked gelatin microgels compared to uncrosslinked gelatin. **E)** Heatmap showing the relative expression of osteogenic genes over time by osteoprogenitor cells seeded on microgels. All gene expressions were normalized to a no-treatment control at day 1, and housekeeping gene Gapdh expression.

**Table 1.** Osteogenic differentiation of progenitor cells seeded on crosslinked gelatin microgels. *Indicates p<0.05

| Tukey multiple comparisons test ($p$-value) | | | | |
| --- | --- | --- | --- | --- |
| Genes | ANOVA | Day 1 vs. Day 3 | Day 1 vs. Day 7 | Day 3 vs. Day 7 |
| *Alp1* | 0.002* | 0.329 | 0.002* | 0.012* |
| *Col1a1* | 0.183 | 0.622 | 0.512 | 0.163 |
| *Runx2* | 0.177 | 0.734 | 0.414 | 0.162 |
| *Bglap* | 0.023* | 0.832 | 0.026* | 0.054 |
| *Spp1* | 0.025* | 0.133 | 0.375 | 0.022* |
| *Ibsp* | 0.004* | 0.741 | 0.004* | 0.010* |
| *Sp7* | 0.071 | 0.345 | 0.403 | 0.061 |

### 3.2. Muscle constructs supported myogenesis and myotubes formation

We created muscle constructs by seeding murine myoblasts (C2C12 cells) in fibrin hydrogels and differentiating them to form myotubes through serum starvation. The differentiation protocol was optimized by culturing myoblasts in tissue culture plastics which showed that five-day serum starvation resulted in well-defined multinucleated myotubes (**Fig.3A**). The differentiation of myoblasts yielded myobundles when seeded in 3D fibrin hydrogels (**Fig.3B**). Further, the temporal gene expression profiles of the myoblasts undergoing differentiation in fibrin hydrogel showed significant changes in the expression levels of prominent skeletal muscle genes (**Fig.3C**). Specifically, the expression of myogenin (*Myog*) continue to be significantly upregulated until day 5 (4-fold increase, *p<0*.*001*), indicating myoblast differentiation to multinucleated myotubes. Likewise, there was a significant upregulation of glucose transporter 4 (*Glut4*), *Mstn, Fndc5*, and interleukin-15 (*Il-15)* over the 5 days, indicating myoblast differentiation in the fibrin constructs. No significant changes in myoblast determination protein 1 (*Myod1*), insulin-like growth factor 1 (*Igf1*), and fibroblast growth factor 2 (*Fgf2*) were noticed by day 5 (See **Table 2** for groupwise comparison).

**Figure 3.**
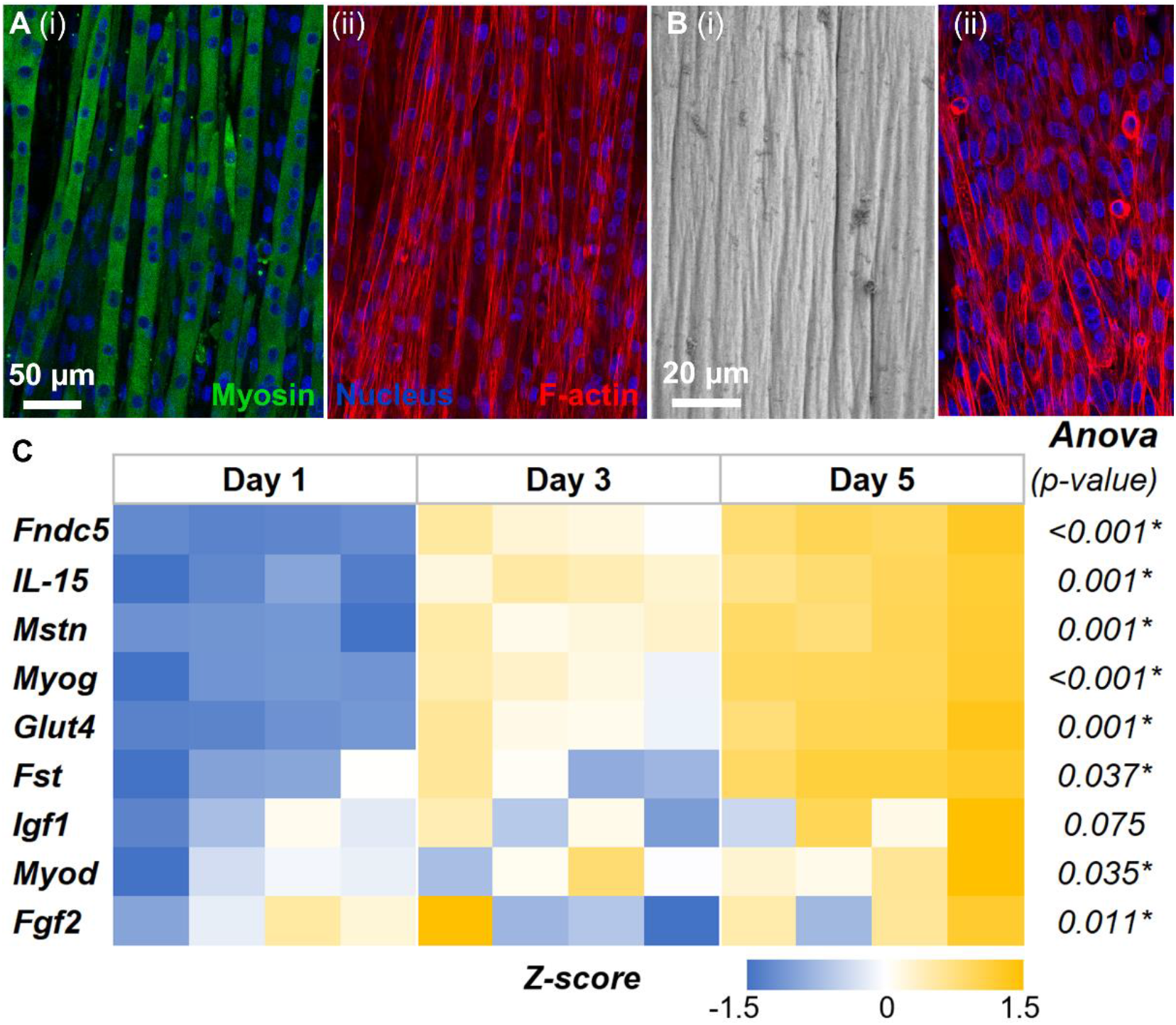
Fabrication and characterization of skeletal muscle constructs. **A)** Confocal image of myoblasts differentiated into multinucleated myotubes expressing myosin in tissue culture plastic. **B)** Scanning electron micrograph of myobundles in 3D fibrin hydrogels. **C)** Temporal gene expression profiles of the myoblasts undergoing differentiation in fibrin hydrogel showed significant changes in the expression levels of prominent skeletal muscle genes. All gene expressions were normalized to a no-treatment control at day 1, and housekeeping gene Ppia expression.

**Table 2.** Myoblast Differentiation. * indicates p<0.05

| Tukey's multiple comparisons test ($p$-value) | | | | |
| --- | --- | --- | --- | --- |
| Genes | ANOVA | Day 1 vs Day 3 | Day 1 vs Day 5 | Day 3 vs Day 5 |
| Fndc5 | <0.001* | 0.006* | <0.001* | 0.012* |
| IL-15 | 0.001* | 0.056 | 0.001* | 0.048* |
| Mstn | 0.001* | 0.038* | 0.001* | 0.064 |
| Myog | <0.001* | 0.005* | <0.001* | 0.002* |
| Glut4 | 0.001* | 0.162 | 0.001* | 0.017* |
| Fst | 0.037* | 0.155 | 0.601 | 0.033* |
| Igf1 | 0.075 | 0.071 | 0.766 | 0.206 |
| Myod | 0.035* | 0.033* | 0.695 | 0.121 |
| Fgf2 | 0.011* | 0.011* | 0.048* | 0.636 |

### 3.3. Exercise training modulates myokine secretion by muscle constructs

We created muscle constructs in 1.2×1.2 cm PDMS wells and subjected them to mechanical stimulation using END and HIIT exercise regimens for 6 hours, and their phenotypes were compared to static controls (**Fig.4A**). A qPCR analysis (**Fig.4B**) showed that *Igf1* and *Il15* were significantly upregulated in both the exercise regimens. However, in END conditions, we also noticed a significant upregulation of *Myog* and downregulation of *Myod1*, indicating differentiation of myoblasts to myotubes. In addition, the osteoinductive myokine irisin (*Fndc5*) was also upregulated in END. There was no significant difference in the expression levels of *Fst, Mstn*, and *Fgf2* under both exercise regimens.

**Figure 4.**
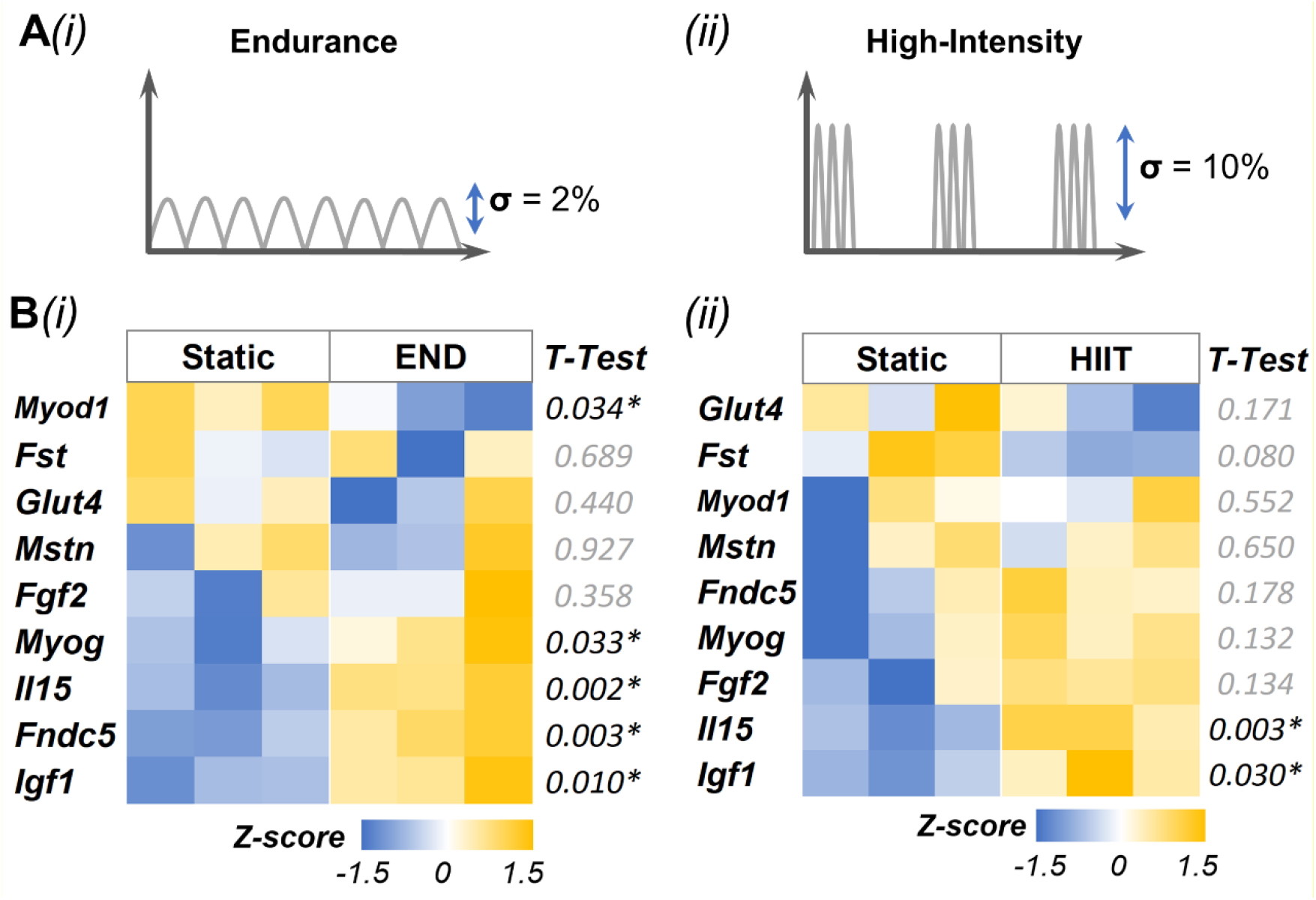
Exercise training modulates muscle phenotype and myokine expression. (**A**) Schematic of exercise regimen i) Endurance (END) and ii) High-intensity interval training (HIIT). (**B**) Gene expression results of i) END and ii) HIIT. All gene expressions were normalized to static control and housekeeping gene Ppia expression.

### 3.4. Exercise training modulates myokine secretion and influences bone phenotype

We studied the effect of exercise training on muscle constructs when they are cocultured with the bone construct. We used a transwell fitted with a 100 µm nylon mesh to separate the bone contructs from the muscle contructs while allowing only soluble factors to diffuse (**Fig.5A**). Then, we independently subjected the muscle constructs to a linear strain. To mimic endurance training (END), the contructs were subjected to 1% strain at 1/30 Hz in bouts of 6 hours for 24 hours with 6 hours intervals (**Fig.5B**). Gene expression analyses showed that myostatin (*Mstn*) was significantly downregulated in cocultured muscle constructs along with *Igf1* and *Glut4* compared to monocultures **(Fig.5C)**. Likewise, *Alp*, a prominent osteogenic marker, was significantly upregulated in the bone constructs cocultured with muscle constructs subjected to END regimen **(Fig.5D)**. But when bone contructs were cocultured with muscle contructs that were not stretched, we noticed significant downregulation of several osteogenic genes (See **Table 3 and 4** for muscle and bone groupwise comparison, respectively).

**Figure 5.**
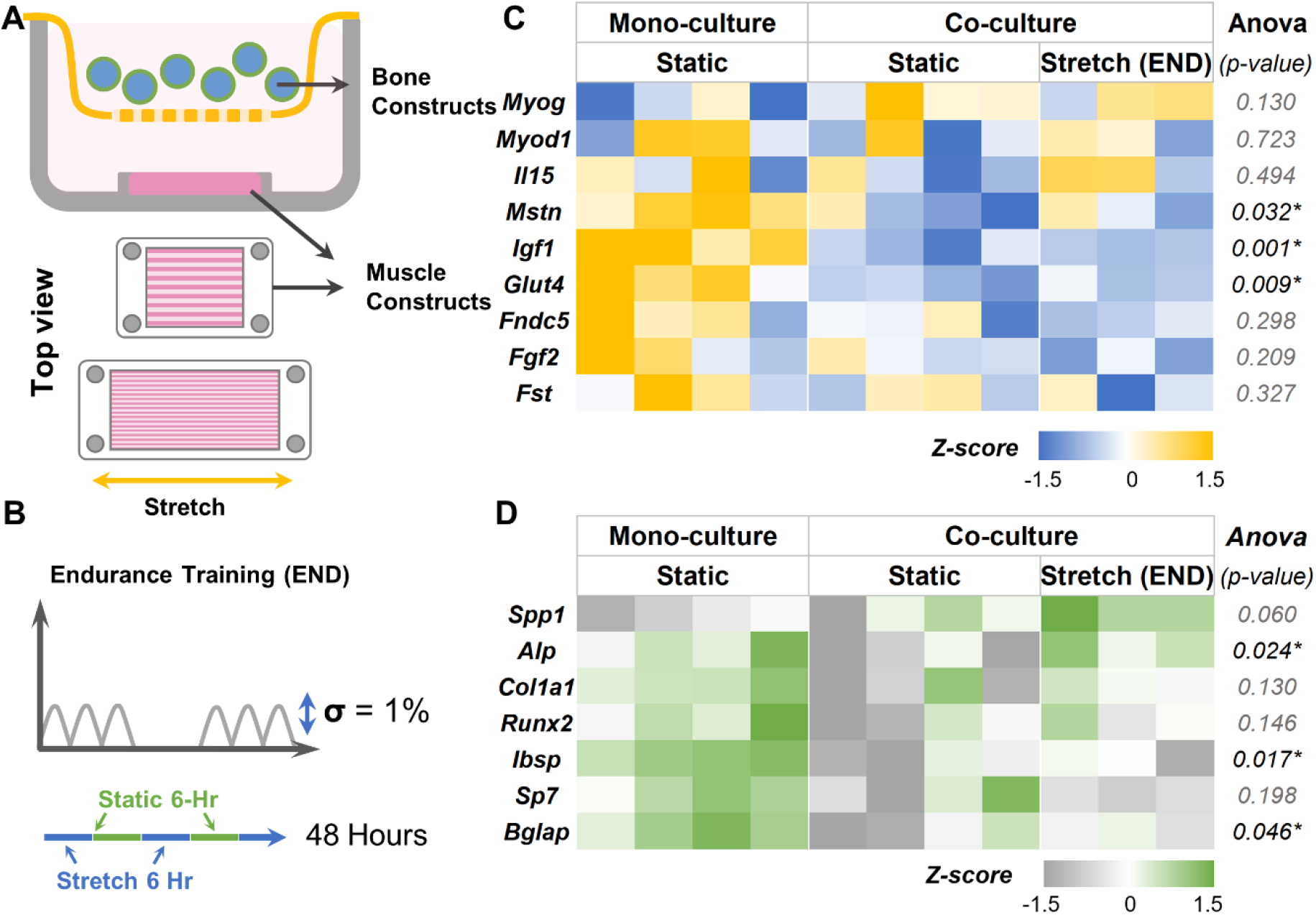
Effects of endurance training on myokine and bone phenotype. **(A)** Schematic showing the setup of the muscle-bone construct cocultures, **(B)** The 48 hours endurance (END) exercise regimen consists of alternating 6-hour stretch (1% strain) at 1/30 Hz and 6 hours rest. **(C)** Heatmaps showing the gene expression of muscle constructs subjected to END regimen when cocultured with bone constructs. **(D)** Gene expression heatmaps of the corresponding bone constructs in the cocultures. All gene expressions were normalized to monoculture controls and housekeeping genes Gapdh (bone) and Ppia (muscle).

**Table 3.**
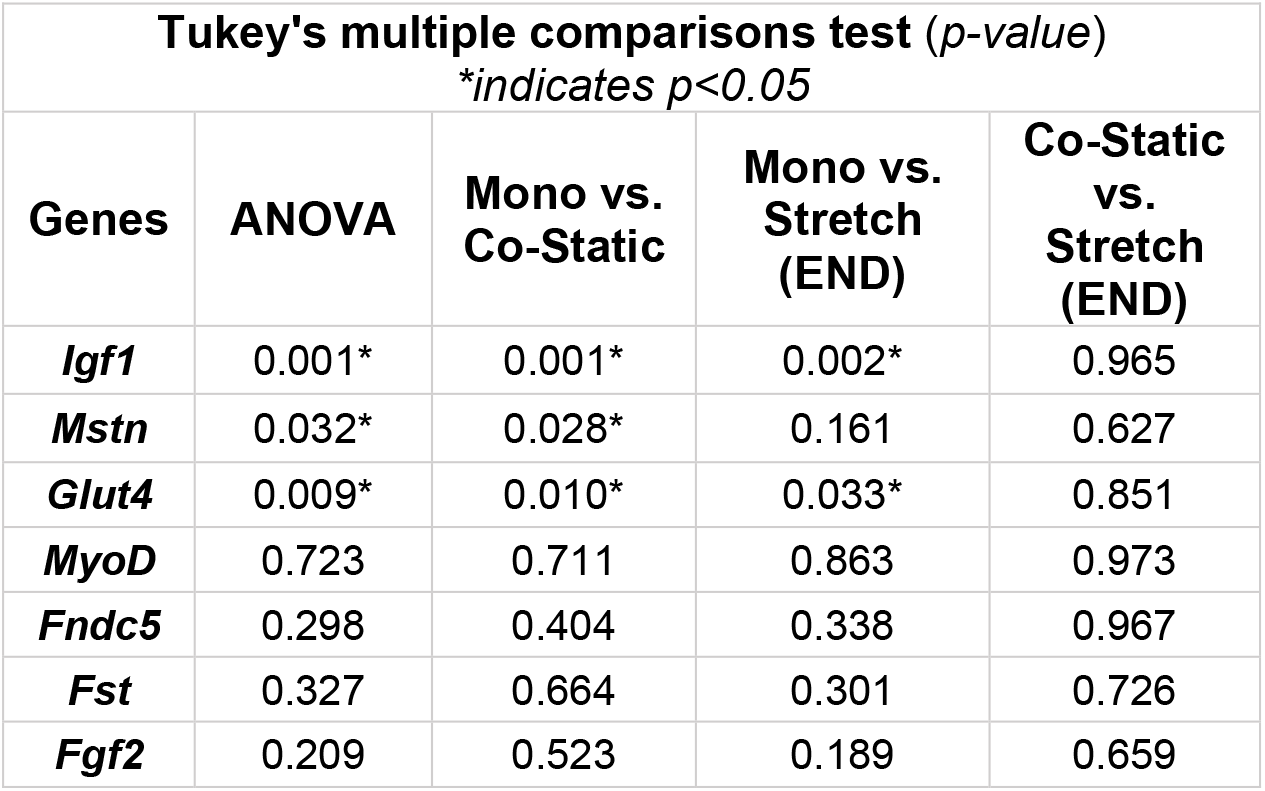
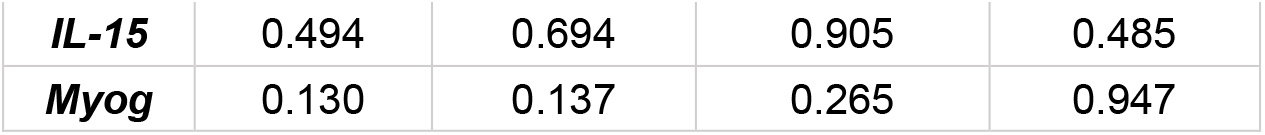
Change in gene expression of muscle constructs subjected to the END.

**Table 4.**
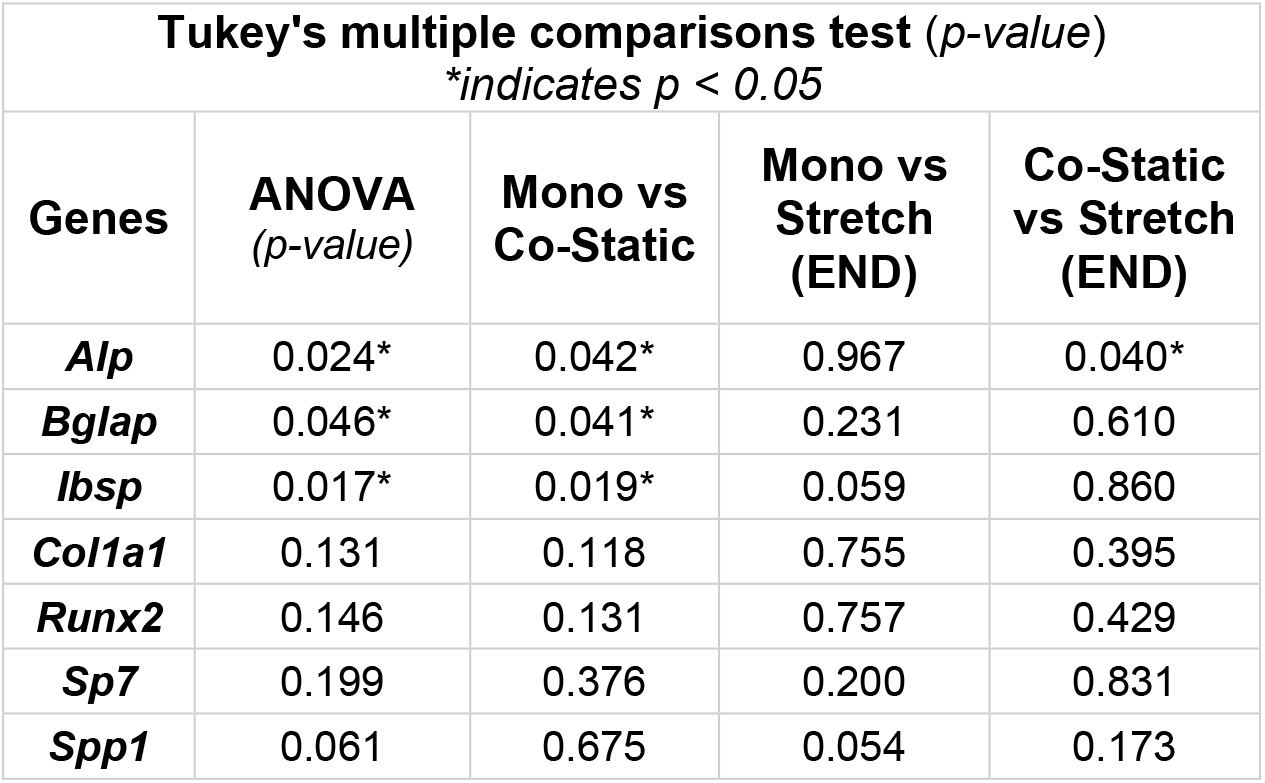
Change in gene expression of bone constructs cocultured with muscle construct (END).

We then investigated the effect of the HIIT exercise regimen **(Fig.6A)** and noticed significant downregulation in *Mstn* in cocultured muscle constructs similar to END exercise training. To mimic HIIT, the constructs were subjected to 10% strain at 1/30 Hz in bouts of 1 hour for 8 hours with 6 hours intervals (**Fig.6B**). Interestingly, expression of *Il15* was significantly increased (*p=0*.*026*) in muscle constructs subjected to HIIT exercise regimen compared to static condition. Further, *Glut4* expression increased, although not statistically significant (*p=0*.*06*), in the cocultured muscle constructs compared to static monocultures (**Fig.6C**, See **Table 5** for groupwise comparison). The altered myokine expression also altered the expression of osteogenic genes in cocultured bone constructs (**Fig.6D**). There was a significant upregulation of genes, including *Spp1* and *Col1a1*, in the cocultured bone constructs compared to static controls (See **Table 6** for groupwise comparison).

**Figure 6.**
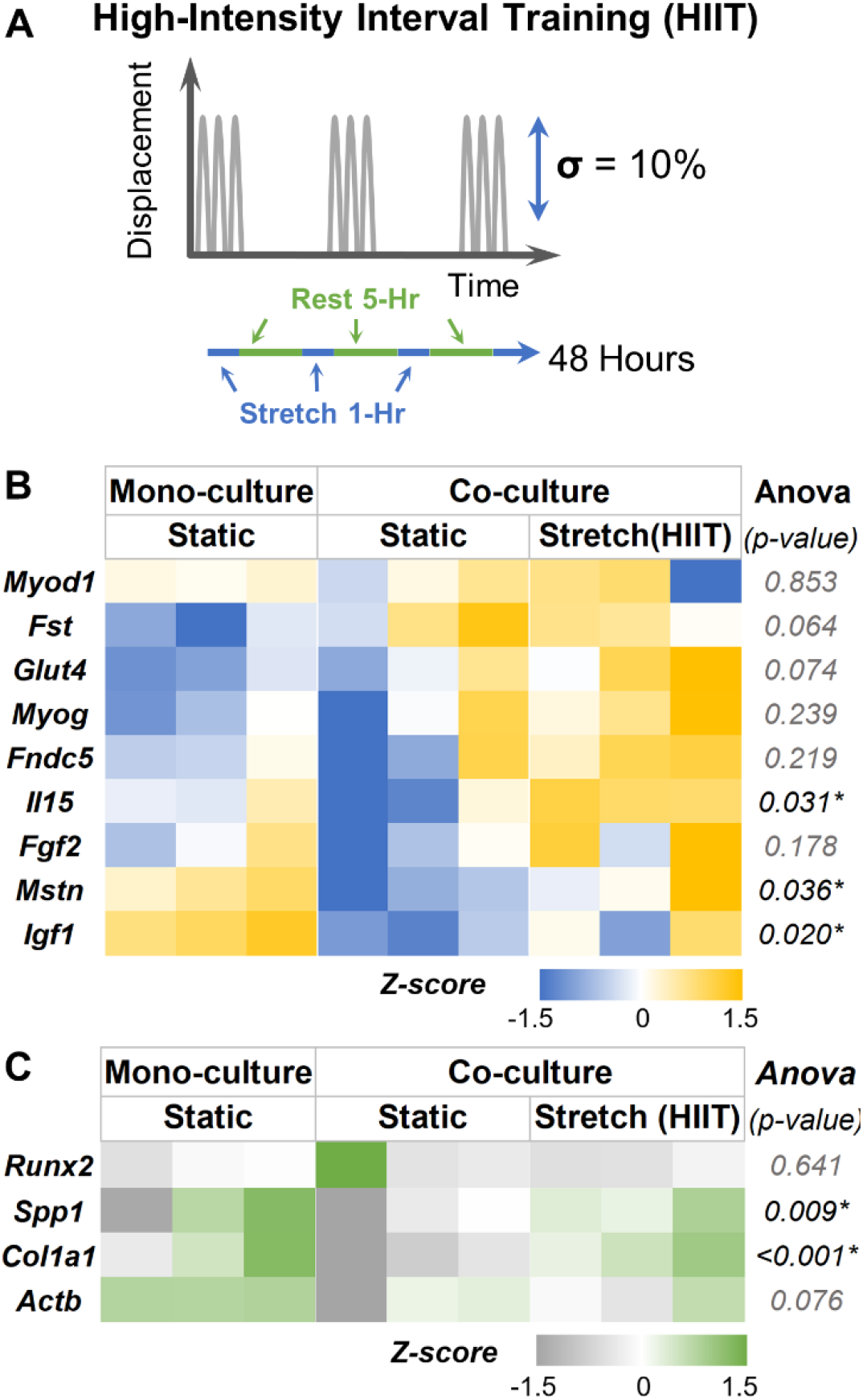
Effects of high-intensity interval (HIIT) training on myokine and bone phenotype. **(A)** The 48 hours high intensity (HIIT) exercise regimen consists of alternating 1-hour stretch (10% strain) at 1/30 Hz and 5 hours of rest. **(B)** Heatmaps showing the gene expression of muscle constructs subjected to HIIT training when cocultured with bone constructs. **(C)** Gene expression heatmaps of the corresponding bone constructs in the cocultures. All gene expressions were normalized to monoculture controls and housekeeping genes Gapdh (bone) and Ppia (muscle).

**Table 5.** Change in gene expression of muscle constructs subjected to the HIIT.

| Tukey's multiple comparisons test (p-value) |  |  |  |  |
| --- | --- | --- | --- | --- |
| * indicates $p < 0.05$ | | | | |
| Genes | ANOVA<br>(p-value) | Mono vs.<br>Co-Static | Mono vs.<br>Stretch<br>(HIIT) | Co-Static<br>vs. Stretch<br>(HIIT) |
| *Igf1* | 0.020* | 0.017* | 0.188 | 0.203 |
| *Mstn* | 0.036* | 0.046* | 0.966 | 0.063 |
| *Il15* | 0.031* | 0.237 | 0.256 | 0.026* |
| *Fst* | 0.064 | 0.079 | 0.103 | 0.976 |
| *Glut4* | 0.074 | 0.545 | 0.065 | 0.263 |
| **Myog** | 0.239 | 0.850 | 0.229 | 0.439 |
| **Myod1** | 0.853 | 0.999 | 0.866 | 0.889 |
| **Fndc5** | 0.219 | 0.940 | 0.343 | 0.228 |
| **Fgf2** | 0.178 | 0.562 | 0.555 | 0.157 |

**Table 6.** Change in gene expression of bone constructs cocultured with muscle construct (HIIT).

| Tukey's multiple comparisons test (*p*-value)<br>*indicates *p* < 0.05 |  |  |  |  |
| --- | --- | --- | --- | --- |
| Genes | ANOVA<br>(*p*-value) | Mono vs.<br>Co-Static | Mono vs.<br>Stretch<br>(HIIT) | Co-Static<br>vs.<br>Stretch<br>(HIIT) |
| **Col1a1** | <0.001* | 0.009* | 0.013* | <0.001* |
| **Spp1** | 0.01* | 0.355 | 0.049* | 0.008* |
| **Actb** | 0.643 | 0.647 | 0.980 | 0.753 |
| **Runx2** | 0.076 | 0.405 | 0.469 | 0.066 |

### 3.5. Myostatin and follistatin influence osteogenesis of progenitor cells

We investigated the impact of the two prominent myokines, myostatin and follistatin, on osteogenic differentiation of preosteoblasts in tissue culture dishes. The preosteoblasts were cultured in both growth and osteogenic conditions. The media was supplemented with myostatin (100 ng/mL), follistatin (100 ng/mL), or a combination of both (100 ng/mL each). In growth conditions, the myostatin treatment significantly downregulated the expression of osteogenic genes *Ibsp, Bglap, Sp7, Alp1*, and *Runx2*, within 48 hours of treatment (**Fig.7A**, See **Table 7** for groupwise comparison). Likewise, when follistatin was supplemented in the growth media, there was a significant downregulation of osteogenic genes *Bglap, Ibsp*, and *Runx2* within 48 hours of treatment. But when compared with the myostatin treatment group, the follistatin treatment significantly upregulated *Ibsp* and *Alp1* expression. When both myostatin and follistatin were supplemented together in the media, there was a significant downregulation of *Bglap, Ibsp, Alp1, and Runx2*. In long-term cultures (7 day, **Fig.7B**, See **Table 8** for groupwise comparison), the myostatin treatment significantly downregulated the expression of *Bglap, Ibsp*, and *Sp7*, similar to the short-term effects. However, follistatin treatment significantly upregulated the osteogenic genes *Bglap, Ibsp*, and *Sp7* in long-term cultures. When both myostatin and follistatin were added, there was a significant down-regulation of *Bgalp, Ibsp*, and *Sp7* compared to the control group. Nevertheless, when compared with the myostatin treatment group, the combinatorial treatment significantly upregulated *Bglap*, and there was no change in the expression levels of other osteogenic genes.

**Figure 7.**
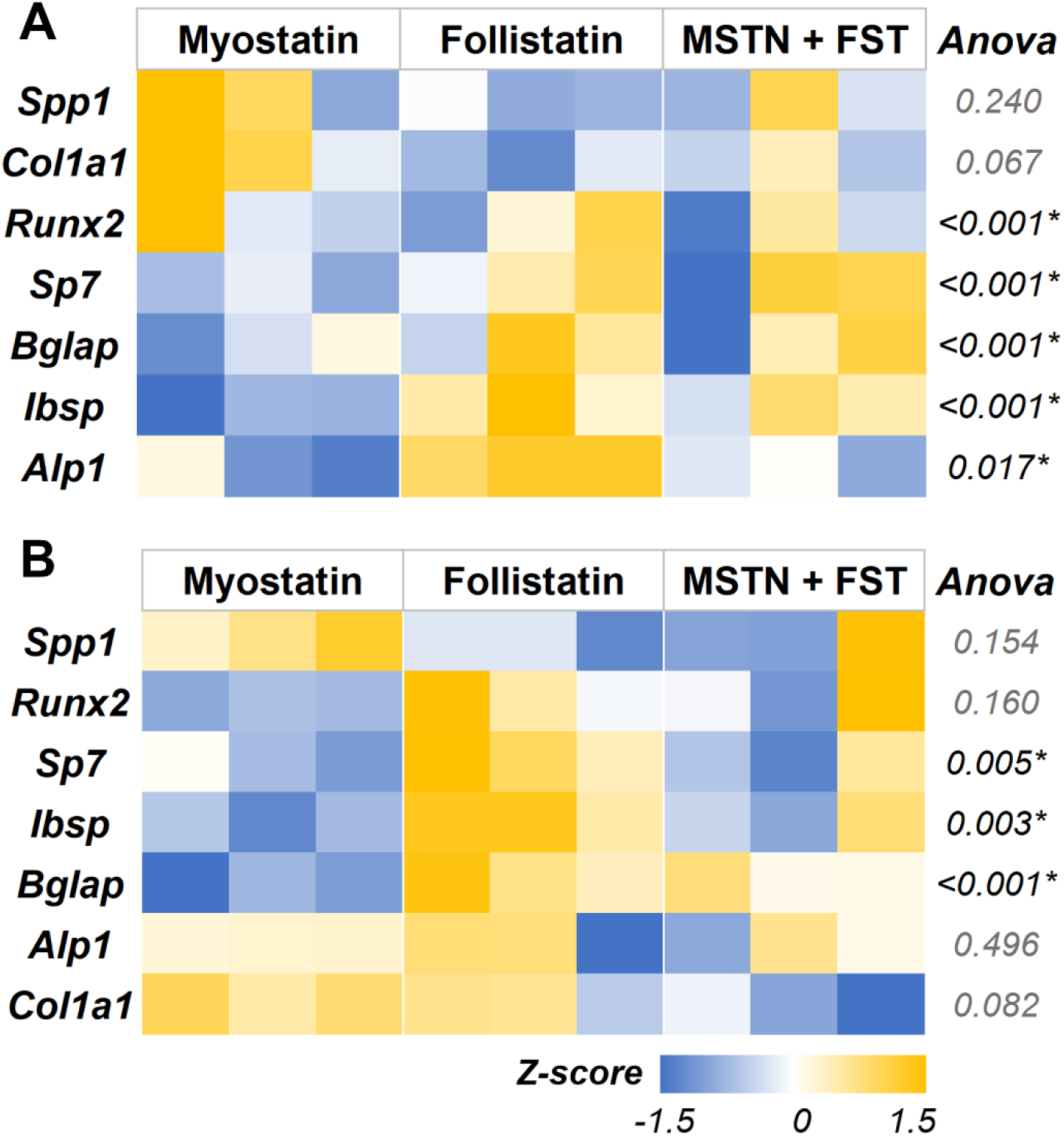
Myostatin treatment negatively regulated prominent osteogenic gene expression. Heatmaps showing the osteogenic gene expression of preosteoblasts cells cultured in growth media supplemented with myostatin, follistatin, or a combination of both for **(A)** 48 hours and **(B)** 7 days. MSTN – Myostatin, FST – Follistatin. All gene expressions were normalized to no-treatment control and housekeeping gene Gapdh expression.

**Table 7.** Gene expression of osteoblasts treated with myostatin and follistatin for 48 Hours. * indicates p<0.05

| Tukey multiple comparisons test (*p*-value) |  |  |  |  |  |  |  |
| --- | --- | --- | --- | --- | --- | --- | --- |
| MSTN – Myostatin, FST – Follistatin |  |  |  |  |  |  |  |
| Genes | ANOVA (*p*-value) | Control vs. MSTN | Control vs. FST | Control vs. MSTN+FST | MSTN vs. FST | MSTN vs. MSTN+FST | FST vs. MSTN+FST |
| *Bglap* | 0.010* | 0.012* | 0.045* | 0.022* | 0.771 | 0.971 | 0.948 |
| *Ibsp* | <0.001* | <0.001* | 0.015* | 0.005* | 0.022* | 0.078 | 0.802 |
| *Alp1* | <0.001* | 0.001* | 0.101 | 0.001* | 0.012* | 0.805 | 0.042* |
| *Sp7* | 0.017* | 0.015* | 0.100 | 0.056 | 0.540 | 0.768 | 0.975 |
| *Runx2* | <0.001* | 0.002* | 0.001* | 0.001* | 0.988 | 0.778 | 0.918 |
| *Spp1* | 0.240 | 0.787 | 0.214 | 0.438 | 0.630 | 0.914 | 0.936 |
| *Col1a1* | 0.066 | 0.997 | 0.125 | 0.374 | 0.095 | 0.295 | 0.838 |
| **Act1** | 0.821 | 0.851 | 0.999 | 0.918 | 0.900 | 0.998 | 0.952 |

**Table 8.** Gene expression of osteoblasts treated with myostatin and follistatin for 7 days. * indicates p < 0.05

| Tukey multiple comparisons test ($p$-value) | | | | | | | |
| --- | --- | --- | --- | --- | --- | --- | --- |
| MSTN – Myostatin, FST – Follistatin |  |  |  |  |  |  |  |
| Genes | ANOVA ($p$-value) | Control vs. MSTN | Control vs. FST | Control vs. MSTN+FST | MSTN vs. FST | MSTN vs. MSTN+FST | FST vs. MSTN+FST |
| **Alp1** | 0.496 | 0.816 | 0.479 | 0.609 | 0.922 | 0.979 | 0.995 |
| **Col1a1** | 0.082 | 0.888 | 0.999 | 0.193 | 0.827 | 0.071 | 0.233 |
| **Runx2** | 0.160 | 0.155 | 0.983 | 0.699 | 0.251 | 0.591 | 0.876 |
| **Bglap** | <0.001* | <0.001* | 0.119 | 0.017* | 0.002* | 0.010* | 0.553 |
| **Spp1** | 0.154 | 0.132 | 0.922 | 0.631 | 0.305 | 0.594 | 0.930 |
| **Ibsp** | 0.003* | 0.005* | 0.914 | 0.038* | 0.012* | 0.487 | 0.097 |
| **Sp7** | 0.005* | 0.009* | 0.529 | 0.013* | 0.067 | 0.995 | 0.094 |

### 3.6. Myostatin knockout in mice altered bone mechanical properties

To study the impact of myostatin on bone mechanical properties, femurs from 7-month-old myostatin knockout mice (GDF8^-/-^) were collected, and three-point bending tests were performed (**Fig.8A**). The whole bone properties, including stiffness, maximum force, work-to-fracture, and post-yield displacement (PDY), and tissue level parameters, including young’s modulus and ultimate stress of femurs from myostatin knockout mice (GDF8^-/-^) were measured and compared to age-matched wildtype mice (**Fig.8B**). After adjusting for body mass, we noticed a significant increase in young’s modulus and decrease in work-to-fracture in myostatin knockout mice (GDF8^-/-^) femurs compared to wildtype (See **Table 9** for raw and adjusted values). No statistical significance was noticed between groups in other measures parameters.

**Figure 8.**
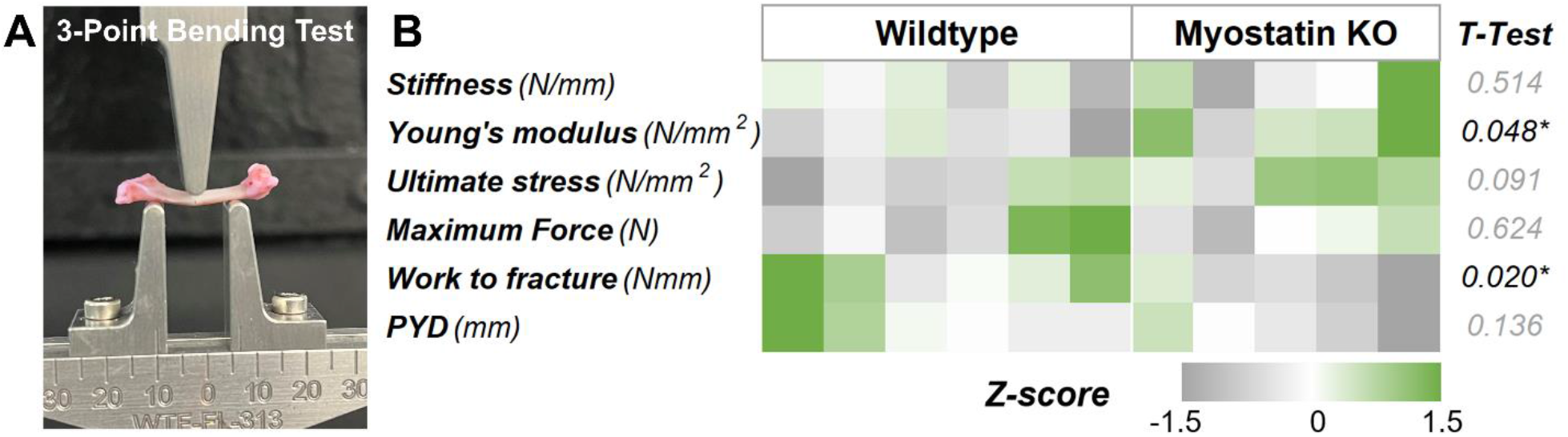
Effect of myostatin knockout on the bone mechanical properties. **(A)** 3-point bending test setting. **(B)** Heatmap comparing the mechanical properties of the femur from myostatin knockout mice (GDF8^-/-^) and wildtype mice.

**Table 9.** Comparison of whole-bone and tissue-level mechanical properties of femur loaded in 3-point bending test. Adult C57BL6 (n = 6; Body mass = 39.5±7 g) and MKO (n = 5; Body mass = 47±4.3 g)

|  | Unadjusted |  | Body mass-adjusted |  |
| --- | --- | --- | --- | --- |
|  | Wildtype | GDF8<sup>-/-</sup> | Wildtype | GDF8<sup>-/-</sup> |
| **Stiffness (N/mm)** | 105.86±15.76 | 115.60±27.74 | 101.29±12.63 | 110.53±27.22 |
|  | 14.88% | 23.99% (0.5) | 12.47% | 24.63% (0.5) |
| **Maximum Force (N)** | 23.147±3.81 | 24.11±0.16 | 23.22±3.81 | 22.31±2.02 |
|  | 16.47% | 11.48% (0.6) | 16.40% | 9.04% (0.6) |
| **Post Yield Displacement (mm)** | 0.439±0.16 | 0.27±0.16 | 0.43±0.16 | 0.27±0.16 |
|  | 37.38% | 58.19 (0.1) | 37.78% | 58.45% (0.1) |
| **Work To Fracture (Nmm)** | 9.872±2.19 | 6.72±2.36 | 10.14±2.12 | 6.28±2.31 |
|  | 22.20% | 35.02 (0.05) | 20.92% | 36.75% (**0.02**) |
| **Young's Modulus (N/mm<sup>2</sup>)** | 1935.08±380.67 | 2236.06±444.76 | 1827.65±309.43 | 2367.44±422.76 |
|  | 19.67% | 19.89% (0.2) | 16.93% | 17.85% (**0.04**) |
| **Ultimate stress (N/mm<sup>2</sup>)** | 58.325±0.00 | 64.40±4.82 | 57.41±7.41 | 64.47±4.82 |
|  | 13.10% | 7.48% (0.1) | 12.89% | 7.47% (0.09) |

Whole-bone mechanical properties were adjusted for body mass using the linear regression method described in the methods. The listed values represent mean ± SD and coefficient of variation (%). P-values are shown in parentheses. Bolded numbers indicate significant differences relative to wildtype control, $p < 0.05$ .

### 3.7. Myostatin knockout altered trabecular and cortical bone properties in mice

To study the impact of myostatin on bone morphological properties, femurs from 7-month-old myostatin knockout mice (GDF8^-/-^) were collected, and bone morphometric analysis was performed through microCT. The trabeculae of a 1.5 mm section of distal metaphysis close to the growth plate were analyzed and compared to age-matched wildtype mice. After adjusting for body mass, we noticed a significant decrease in the trabecular separation (Tb.Sp), trabecular pattern (Tb.Pf), and structural model index (SMI) in femurs from myostatin knockout (GDF8^-/-^) mice compared to wildtype (**Fig.9A**, See **Table.10** for raw and adjusted values). The 3D surface rendering showed more connected and denser trabeculae in the myostatin knockout (GDF8^-/-^) mice compared to wildtype **(Fig.9B)**.

**Figure 9.**
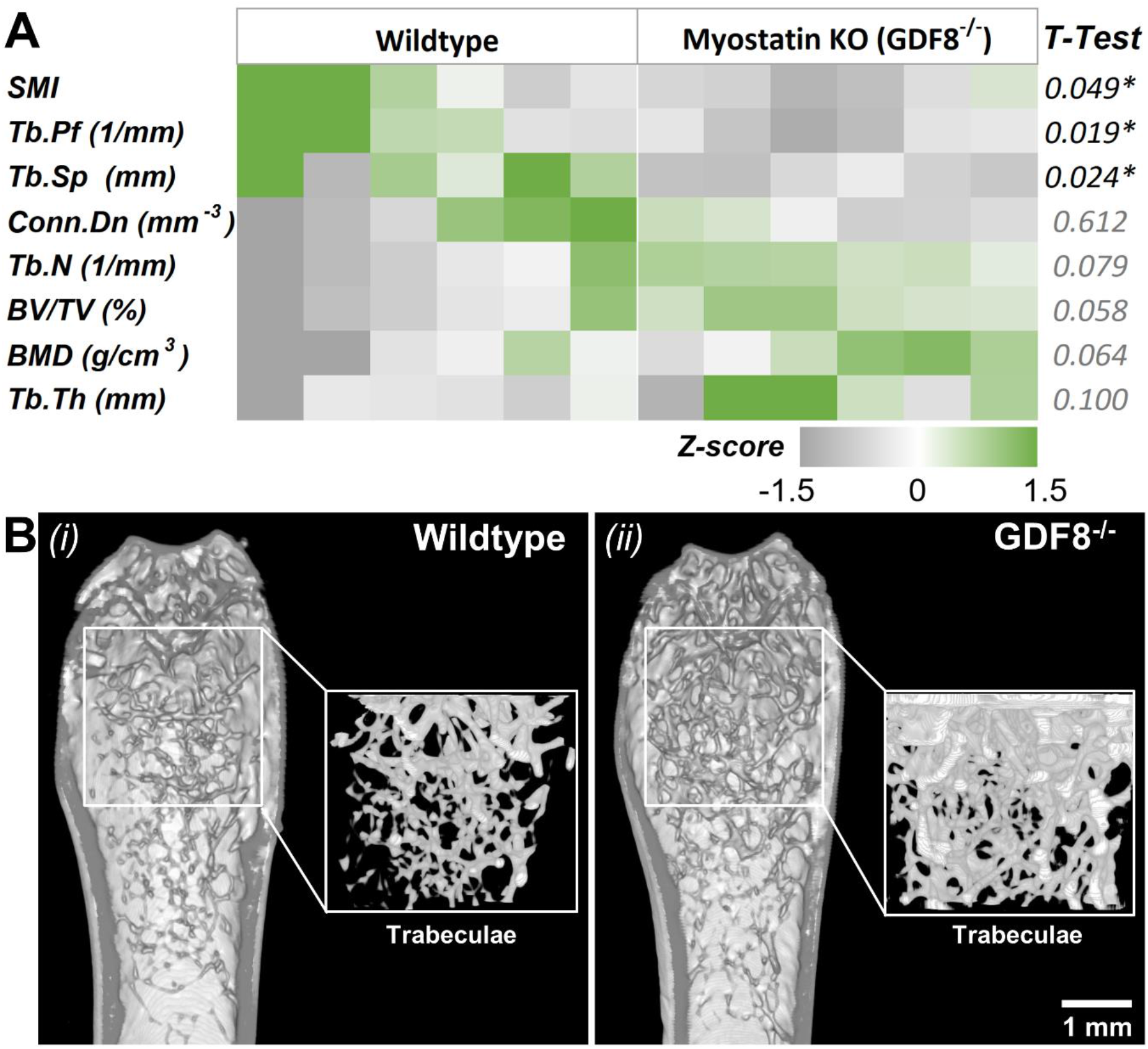
Effect of myostatin knockout on the morphometric properties of trabeculae. **(A)** Heatmap comparing the bone morphometric parameters of the distal femur trabeculae of myostatin knockout (GDF8^-/-^) and wildtype mice. (B) Sagittal cross-sectional of distal metaphysis showing the trabeculae of (i) Wildtype and (ii) GDF8^-/-^ mice. Insets show the 3D surface rendering of the trabeculae alone in the distal metaphysis area. SMI - Structural model index, Tb.Pf - Trabecular pattern, Tb.Sp - Trabecular separation, Conn.Dn - Connectivity density, Tb. N - Trabecular number, BV/TV - Percent bone volume, BMD - Bone mineral density, Tb.Th - Trabecular thickness.

**Table 10.** Comparison of unadjusted and body-mass adjusted values for trabecular bone structural properties of femurs. Adult wildtype C57BL6 (WT, n = 6; Body mass = 39.5±7 g) and myostatin knockout (MKO, n = 5; Body mass = 47±4.3 g)

|  |  | Unadjusted |  | Body mass-adjusted |  |
| --- | --- | --- | --- | --- | --- |
|  |  | WT | MKO | WT | MKO |
| Percent bone volume (%) | BV/TV | 14.346±4.158 | 16.299±0.898 | 13.078±2.987 | 16.060±0.808 |
|  |  | 28.98% | 5.51% (0.3) | 22.84% | 5.03% (0.05) |
| **Trabecular pattern factor (1/mm)** | **Tb.Pf** | 16.408±2.473<br>15.07% | 14.397±1.233<br>8.56% (0.11) | 16.952±2.139<br>12.61% | 14.025±1.072<br>7.64% (**0.01**) |
| **Trabecular thickness (mm)** | **Tb.Th** | 0.112±0.002<br>2.17% | 0.114±0.005<br>4.07% (0.27) | 0.111±0.002<br>2.00% | 0.115±0.004<br>3.78% (0.09) |
| **Trabecular number (1/mm)** | **Tb.N** | 1.279±0.350<br>27.35% | 1.427±0.072<br>5.01% (0.35) | 1.171±0.249<br>21.23% | 1.395±0.047<br>3.40% (0.07) |
| **Trabecular separation (mm)** | **Tb.Sp** | 0.369±0.095<br>25.72% | 0.312±0.031<br>9.88% (0.21) | 0.402±0.058<br>14.45% | 0.327±0.018<br>5.47% (**0.02**) |
| **Structure model index** | **SMI** | 2.640±0.224<br>8.48% | 2.497±0.107<br>4.29% (0.20) | 2.684±0.200<br>7.46% | 2.470±0.098<br>3.94% (**0.04**) |
| **Connectivity density (1/mm<sup>3</sup>)** | **Conn.Dn** | 15.984±8.592<br>53.75% | 13.445±3.113<br>23.16% (0.53) | 14.338±7.727<br>53.89% | 12.552±2.749<br>21.90% (0.6) |
| **Bone mineral density (g/cm<sup>3</sup>)** | **BMD** | 0.162±0.019<br>11.67% | 0.168±0.008<br>4.45% (0.50) | 0.156±0.012<br>7.61% | 0.168±0.007<br>4.46% (0.06) |

Trabecular structural properties were adjusted for body mass using the linear regression method described in the methods. The listed values represent mean±SD and coefficient of variation (%). *p* values are shown in parentheses. Bolded numbers indicate significant differences relative to WT, *p*<0.05.

For comparing the cortical bone properties, a 2 mm section of mid diaphysis was analyzed and compared to age-matched wildtype mice. We found that the bone volume fraction (BV/TV) was significantly reduced, and the total cortical porosity percentage was significantly higher in femur femurs from myostatin knockout (GDF8^-/-^) mice compared to wildtype (**Fig.10A**, See **Table.9** for raw and adjusted values). In addition, the 3D surface rendering showed slightly thinner cortical bone in the myostatin knockout (GDF8^-/-^) mice compared to wildtype **(Fig.10B)**.

**Figure 10.**
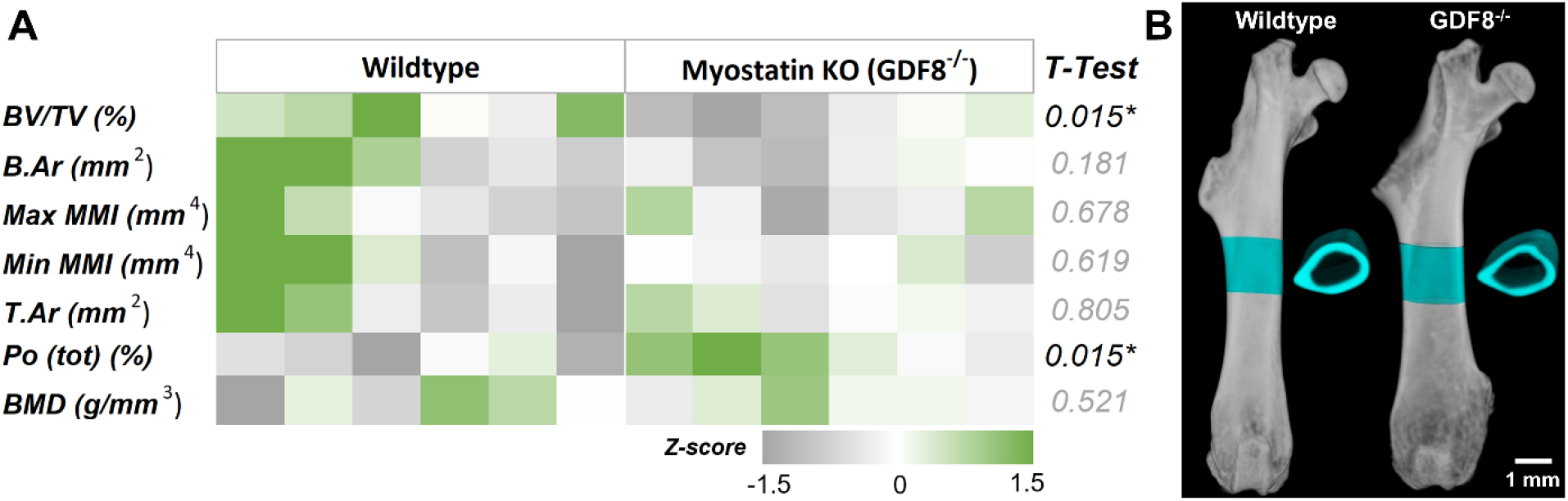
Effect of myostatin knockout on the morphometric properties of cortical bone. **(A)** Heatmap comparing the bone morphometric parameters of the cortical bone in mid-diaphysis of myostatin knockout (GDF8^-/-^) and wildtype mice. BV/TV - Percent bone volume (bone volume/Tissue volume), B.Ar - Mean total cross-sectional bone area, Max MMI - Average principal moment of inertia (max), Min MMI - Average principal moment of inertia (min), T.Ar - Mean total cross-sectional tissue area, Po(tot) -Total porosity, BMD - Bone mineral density. **(B)** 3D surface rendering of the whole femur and the crosssection of a 2 mm cortical bone in the mid-diaphysis.

**Table 9.** Comparison of unadjusted and body-mass adjusted values for cortical bone structural properties of femurs. Adult wildtype C57BL6 (WT, n = 6; Body mass = 39.5±7 g) and myostatin knockout (MKO, n = 5; Body mass = 47±4.3 g)

|  |  | Unadjusted |  | Body mass-adjusted |  |
| --- | --- | --- | --- | --- | --- |
|  |  | WT | MKO | WT | MKO |
| **Bone mineral density** | **BMD** | 1.267±0.060 | 1.272±0.019 | 1.254±0.052 | 1.269±0.018 |
|  |  | 4.72% | 1.48% (0.83) | 4.14% | 1.42% (0.52) |
| **Total porosity (percent)** | **Po(tot)** | 54.729±2.023 | 57.893±1.980 | 55.224±1.679 | 58.265±1.884 |
|  |  | 3.70% | 3.42% (**0.02**) | 3.04% | 3.23% (**0.01**) |
| **Percent bone volume** | **BV/TV** | 45.271±2.023 | 42.107±1.980 | 44.776±1.679 | 41.735±1.884 |
|  |  | 4.47% | 4.70% (**0.02**) | 3.75% | 4.51% (**0.01**) |
| **Mean total crosssectional tissue area** | **T.Ar** | 2.176±0.221 | 2.223±0.070 | 2.225±0.192 | 2.246±0.059 |
|  |  | 10.17% | 3.14% (0.63) | 8.63% | 2.64% (0.80) |
| **Mean total crosssectional bone area** | **B.Ar** | 0.991±0.095 | 0.943±0.039 | 1.003±0.090 | 0.943±0.039 |
|  |  | 9.57% | 4.13% (0.29) | 9.00% | 4.13% (0.18) |
| **Average principal moment of inertia (max)** | **Av.MMI(max)** | 0.375±0.060 | 0.372±0.036 | 0.390±0.050 | 0.379±0.034 |
|  |  | 15.93% | 9.68% (0.91) | 12.94% | 9.03% (0.67) |
| **Average principal moment of inertia (min)** | **Av.MMI(min)** | 0.189±0.044 | 0.183±0.013 | 0.195±0.042 | 0.186±0.012 |
|  |  | 23.26% | 7.14% (0.75) | 21.46% | 6.60% (0.61) |

Cortical structural properties were adjusted for body mass using the linear regression method described in the text. The listed values represent mean ± SD and coefficient of variation (%). $p$ values are shown in parentheses. Bolded numbers indicate significant differences relative to WT, $p < 0.05$ .

## 4. Discussion

Uncoupling the effects of biophysical and biochemical stimuli on the adaptive response of bone during exercise training is challenging. We addressed this limitation by designing an *in-vitro* coculture system that allows independent mechanical manipulation of bone and muscle constructs. The muscle constructs were created in a stretchable PDMS well by seeded murine myoblasts in fibrin hydrogels that formed multinucleated myotubes upon serum starvation. The reduction in *Myod* expression and the increase in *Myog* levels are indicators of myoblast differentiation into myotube^43^. Early expression of *Myod*is essential for myoblast proliferation and induction of differentiation, whereas the late expression of *Myog* is vital for the terminal differentiation and fusion of myoblast into mature muscle fibers^44^. Fibrin hydrogels allowed modulating mechanical properties, including stiffness and pore size, and favored myogenesis by supporting myoblast proliferation and myotube formation^45^. On the other hand, the crosslinked gelatin microgels supported the osteogenic differentiation of progenitor cells. The physical properties, including the curvature, stiffness, and surface characteristics of the microgels, contributed significantly to its osteoconductive property^34, 46, 47^. Using these constructs we evaluated the effects of mechanical strain on myokine secretion and its impact on bone metabolism decoupled from physical stimuli.

Exercise training has been shown to have protective effects on muscle and bone biology. Exercise training can activate resident myogenic stem cells, aka satellite cells, to produce hepatocyte growth factor (HGF) that can promote myogenesis, muscle repair, and regeneration^48^. In addition, exercise training influences the cellular metabolism of muscles, including glucose uptake and phosphorylation of anabolic targets^49^. Our studies show that both HIIT and END exercise regimens increase the expression levels of *Igf1* and *Il15*. However, END exercise training seems to contribute significantly to the differentiation of myoblasts, as evident from the upregulation of *Myog* and downregulation of *Myod1*. In addition, the osteoinductive myokine irisin (*Fndc5*) was significantly upregulated in END, indicating that END regimen may be more suitable for increasing bone quality and density^50^ while contributing to muscle hypertrophy^50, 51^. But when cocultured with bone constructs, the effect of the exercise regimens changed markedly. The modular nature of the constructs enabled coculturing the muscle constructs with bone contructs in a biomechanically decoupled fashion. Under static conditions, there was an upregulation of *Myog* and *Fst* in muscle constructs and *Spp1* (Osteopontin) in bone constructs. Osteopontin is a matricellular protein that promotes proliferation and calcification in osteoblasts and mediates bone metabolism^52^. Further, *Mstn* expression was significantly downregulated in muscle constructs cocultured with bone constructs compared to monocultures. Myostatin can bind to activin receptor IIB (ActRIIB) in osteogenic cells and negatively regulate osteoblast and osteoclast gene expression^53^. The role of myostatin in skeletal muscle is redundant with another protein of the TGFβ family, activin A. But inhibiting myostatin activity in muscles improves muscle health and promotes muscle hypertrophy^54^.

Follistatin, an endogenous inhibitor of myostatin, is also shown to promote muscle growth^32^. Hence, the secreted factors from muscle and bone constructs influence each other in the absence of biomechanical stimuli. Although, the factors that are secreted by osteoblasts are unclear in our study.

When muscle constructs were independently subjected to mechanical stimulation, we saw changes to both muscle and bone phenotype depending on the exercise regimen. Notably, *Igf1* was significantly downregulated in cocultured muscle constructs subjected to END but not in HIIT. Consequently, the insulin/IGF1-regulated glucose transporter, *Glut4*, was significantly downregulated in cocultured muscle construct under END regimen but upregulated in the HIIT. Other studies have also shown that endurance training can reduce muscle glucose uptake due to reduced *GLUT4* translocation^55, 56^, while acute bouts of exercise can increase their translocation in an insulin-independent manner^57^. The cocultured bone constructs also showed variable phenotype changes when the muscle constructs underwent exercise training. Early osteogenic marker, *Alp*, was significantly upregulated in END regimen, while late-stage osteogenic markers, *Col1a1* and *Spp1*, were significantly upregulated in the HIIT regimen.

Hence, the myokines secreted under the HIIT regimen seem to promote a healthier bone phenotype than END training. It is possible that the degree to which *Mstn* is expressed or suppressed in muscle constructs primarily determines the beneficial effect on bone phenotype. Studies show that decoy receptors of myostatin enhance the osteogenesis of progenitors^58^ and consequently increase bone mass and density^59^. Hence, we analyzed the role of myostatin and follistatin on the phenotype of the bone constructs isolated from other myokines. Our *in vitro* studies confirmed the adverse effects of myostatin on bone constructs by significantly reducing osteogenic gene expressions of the osteoprogenitor cells. When we treated the bone constructs with myostatin and follistatin, we noticed that myostatin’s inhibitory effects on osteogenic differentiation significantly ameliorated. More mechanistic studies using selective inhibitors of osteokines and myokines could further clarify their specific roles in physiology and pathologies.

Finally, using myostatin knockout (GDF8^-/-^) mice, we analyzed the role of myostatin on bone structure and function. Myostatin knockout mice are generally muscular compared to age-matched wildtype, but their body mass is not significantly different (*p = 0*.*06*). In addition, the measured whole-bone stiffness was not significantly different between the groups; however, the young’s modulus, a measure of the “tissue level stiffness^42^,” was significantly higher in GDF8^-/-^ femurs than in the wildtype after adjusting for body mass. Likewise, the work-to-fracture, an integrative measure of a structure’s overall resistance to failure^42^, is lower in GDF8^-/-^ compared to wildtype. These differences were reflected in cortical and trabecular microarchitecture analyzed through microCT. Although the normalized trabecular bone volume is not significantly different (*p= 0*.*058*) in GDF8^-/-^ femurs, they exhibited significantly lower trabecular pattern factor and trabecular separation compared to wildtype. Similarly, the GDF8^-/-^ femurs exhibited lower cortical bone volume and higher porosity, indicating signs of bone loss. Hence, our data suggest that, in the absence of comorbidities, myostatin knockout (GDF8^-/-^), although enhanced muscle mass, moderately influences bone phenotype in adult mice. However, other studies in mouse models of osteoporosis, osteoarthritis, osteogenesis imperfecta, muscular dystrophy, and diabetes show that pharmacological inhibition of myostatin improves muscle and bone properties to a greater extend^60^. Myostatin inhibition also ameliorates obesity-induced bone loss and trabecular architecture deterioration in rats^61^. The variation could be partially due to the redundant role of myostatin (GDF8) with another TGF-β family member, growth differentiation factor 11 (GDF11), in skeletal patterning but not in regulating muscle mass^62^. Further pharmacological inhibition of myostatin usually utilizes decoy receptors (AcvrIIB-Fc) or neutralizing antibodies that do not differentiate between GDF8 and GDF11^63^, as mature GDF8 shares 90% sequence homology with GDF11 and are highly identical in vertebrate model animals^64^. A study that used myostatin-specific antibodies (Mstn-mAb) reported no significant change in bone mass^63^. Likewise, another study reported that myostatin deficiency enhanced osteoblast differentiation in a load-dependent manner. However, in the absence of loading, it did not protect against bone loss in a mouse model^65^.

Hence, the myostatin-deficient mouse model is suitable for studying changes in muscle phenotype. However, it is challenging to demarcate their effects on bone independent of confounding variables associated with mechanical loading. On the other hand, our *in-vitro* coculture system was able to elucidate the effects of different exercise training regimens on myokine secretion and its impact on bone metabolism in a diabetic environment. END exercise training seems to contribute more significantly to the differentiation of myoblasts than HIIT. But when cocultured with bone constructs, the myokine secretion in the HIIT is more likely to enhance muscle phenotype. Likewise, the myokines secreted under the HIIT regimen promote a healthier bone phenotype than END training. Together, our *in vitro* studies suggest that the type of exercise influences muscle phenotype and the secreted biochemical signals from muscle and bone constructs can influence each other without biomechanical stimuli.

Overall, our coculture system allowed orthogonal manipulation of mechanical strain on muscle constructs while facilitating biochemical crosstalk between bone and muscle constructs. In addition, our system is innovative as it provides an individualized microenvironment and allows decoupled biomechanical manipulation, which is unachievable using traditional models. In the long-term, these in-vitro systems can help identify molecular targets and develop engineered therapies for diabetic bone disease.

## 6. Acknowledgments

Research reported in this publication was supported in part by the National Institute of Arthritis and Musculoskeletal and Skin Diseases (NIAMS) Award Number R21AR078447, National Institute of General Medical Sciences (NIGMS) of the National Institutes of Health under Award Numbers P20GM130456 and P20GM103436-20 (KY IDeA Networks of Biomedical Research Excellence), Igniting Research Collaborations program funded by the University of Kentucky Vice President for Research and College Deans, and Orthopedic Trauma Association (OTA, Grant Number: 6889). The content is solely the responsibility of the authors and does not necessarily represent the official views of the National Institutes of Health or other grant funding agencies. The myostatin knockout mice were a kind donation from Regeneron Pharmaceuticals.

## 7. Supplementary information

Supplement information (DOCX)

**Supplementary Figure 1.**
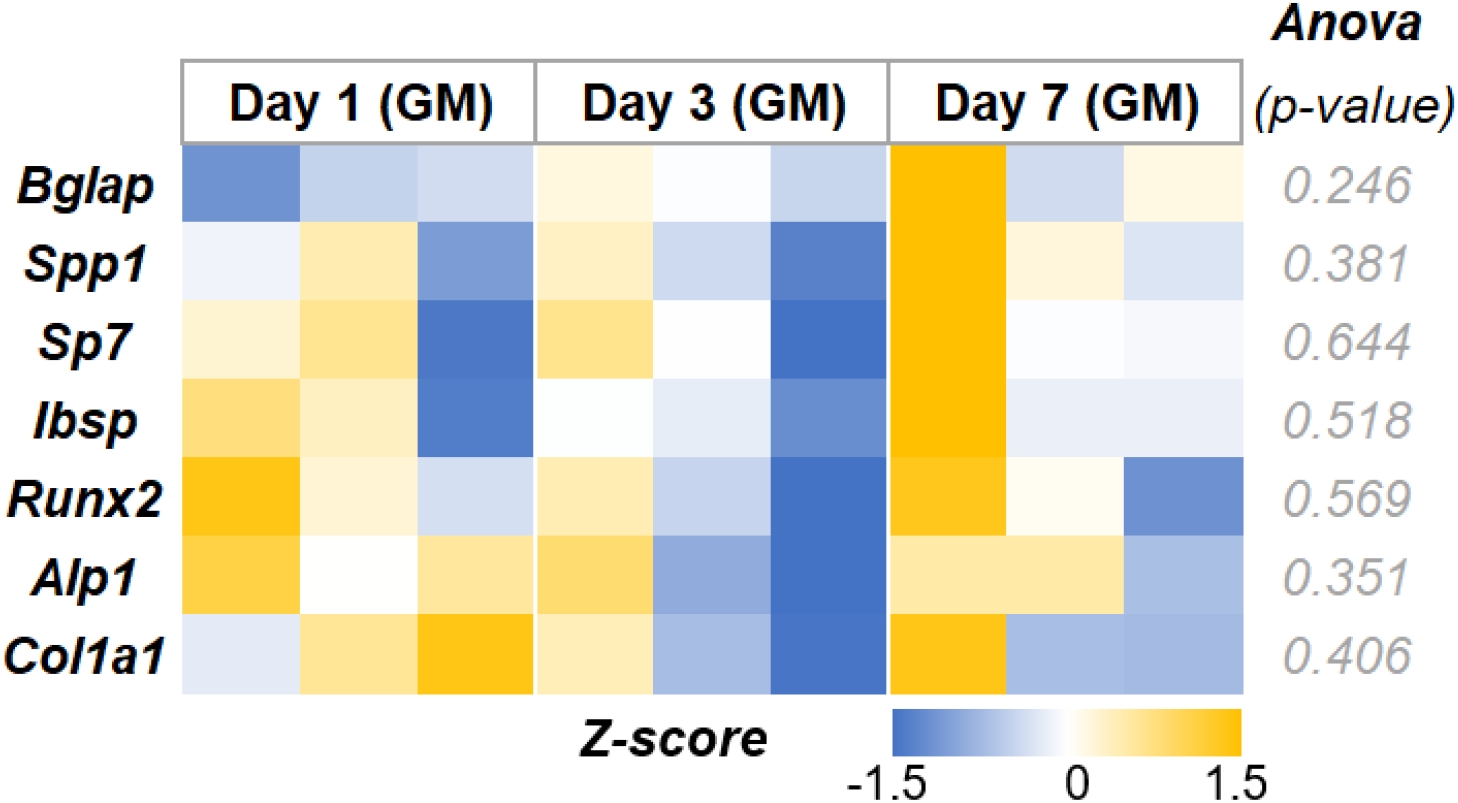
Heatmaps depicting the osteogenic gene expression of cells seeded on crosslinked gelatin microgels and cultured in growth media.

**Supplementary Figure 2.**
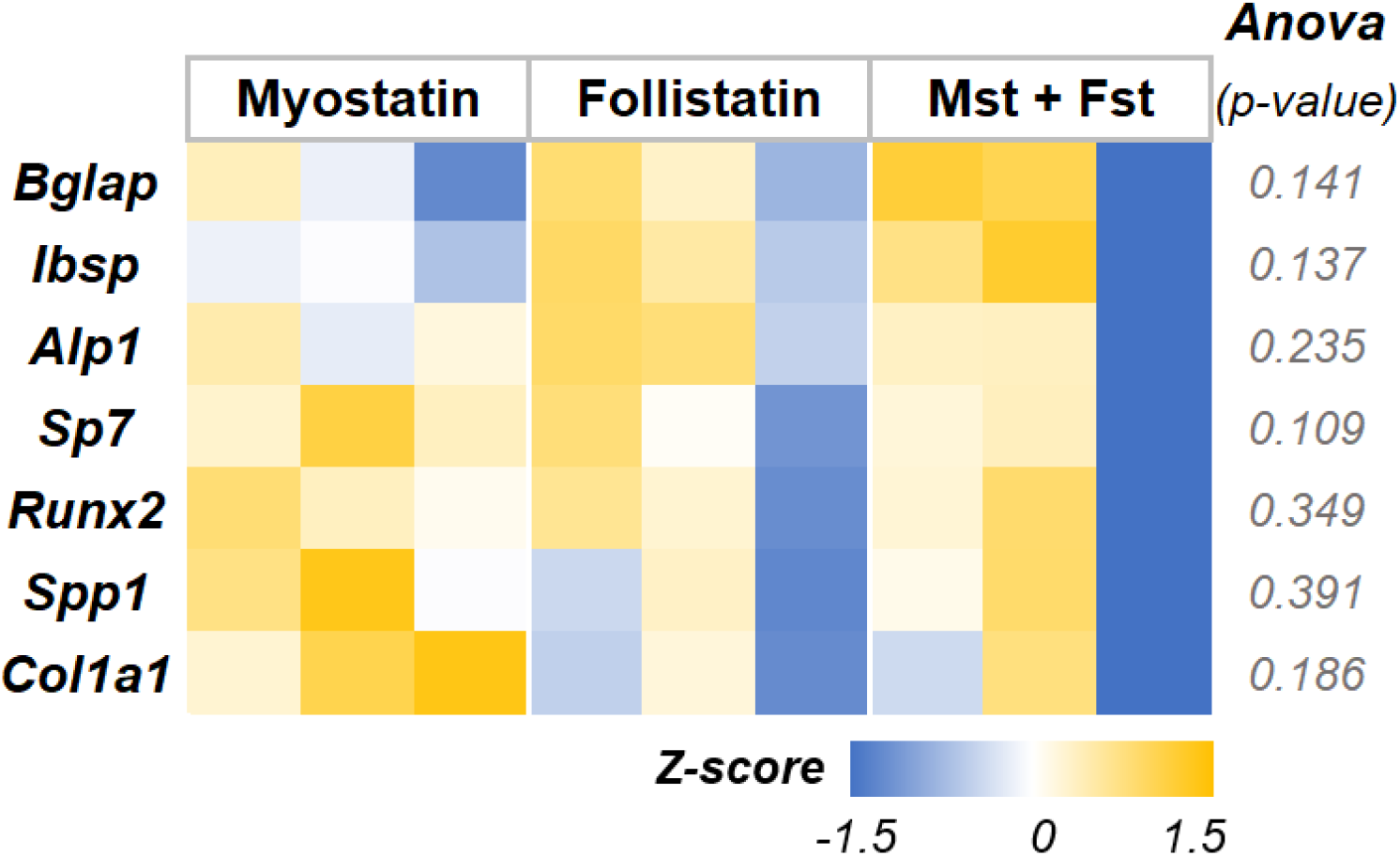
Heatmaps depicting the osteogenic gene expression of cells cultured in osteogenic media supplemented with myostatin, follistatin or a combination of both.

